# White matter microstructural properties in bipolar disorder and its relationship to the spatial distribution of lithium in the brain

**DOI:** 10.1101/346528

**Authors:** Joe Necus, Nishant Sinha, Fiona Elizabeth Smith, Peter Edward Thelwall, Carly Jay Flowers, Peter Neal Taylor, Andrew Matthew Blamire, David Andrew Cousins, Yujiang Wang

**Affiliations:** Institute of Neuroscience, Newcastle University, Newcastle upon Tyne, NE1 7RU, United Kingdom; Interdisciplinary Computing and Complex BioSystems (ICOS), School of Computing Science, Newcastle University, Newcastle upon Tyne, NE1 5TG, United Kingdom; Institute of Cellular Medicine, Newcastle University, Newcastle upon Tyne, NE1 7RU, United Kingdom; Newcastle Magnetic Resonance Centre, Newcastle University, Campus for Ageing and Vitality, Newcastle upon Tyne, NE4 5PL, United Kingdom; Institute of Neurology, University College London, London, WC1N 3BG, United Kingdom.

**Keywords:** Bipolar Disorder, Lithium, Magnetic Resonance Imaging, Diffusion Imaging, White Matter, Fractional Anisotropy

## Abstract

**Background:** Lithium treatment is associated with an increase in magnetic resonance imaging derived measures of white matter integrity, but the relationship between the spatial distribution of brain lithium and white matter integrity is unknown.

**Methods:** Euthymic patients with bipolar disorder receiving lithium treatment (n=12) and those on other medications but naïve to lithium (n=17) underwent diffusion imaging alongside matched healthy controls (n=16). Generalised fractional anisotropy (gFA) within white matter was compared between groups using a standard space white matter atlas. Lithium-treated patients also underwent novel multinuclear 3D lithium magnetic resonance imaging (^7^Li-MRI) to determine relative lithium concentration across the brain. The relationship between ^7^Li-MRI signal intensity and gFA was investigated at the resolution of the ^7^Li-MRI sequence in native space.

**Results:** The lithium-treated bipolar disorder and healthy control groups had higher mean gFA in white matter than the bipolar disorder group treated with other medications but naïve to lithium (t = 2.5, *p* < 0.05; t = 2.7, *p* < 0.03, respectively). No differences in gFA were found between patients taking lithium and healthy controls (t = 0.02, *p* = 1). These effects were seen consistently across most regions in the white matter atlas. In the lithium-treated group, a significant effect of the ^7^Li-MRI signal in predicting the gFA (*p* < 0.01) was identified in voxels containing over 50% white matter.

**Conclusions:** Lithium treatment of bipolar disorder is associated with higher gFA throughout brain white matter, and the spatial distribution of lithium is also positively associated with white matter gFA.

## INTRODUCTION

Bipolar disorder is a major mental illness affecting 1% of the world’s population. Although numerous treatments are available, lithium retains a key position in major treatment guidelines, effectively treating and preventing illness episodes and reducing suicidal behaviour (Tondo, Vazquez, & Baldessarini, 2014). The mechanisms by which lithium achieves these effects are not fully understood and we lack detailed information about its tissue level distribution and how this relates to its actions in the brain.

Various neuroimaging techniques have been used to identify structural and functional changes that occur within the brain during treatment with lithium. For instance, cross-sectional structural MRI studies have revealed that treatment with lithium is associated with increased grey matter volume and cortical thickness (Hafeman, Chang, Garrett, Sanders, & Phillips, 2012; Hibar et al., 2017; Hibar et al., 2016; Moore, Bebchuk, Wilds, Chen, & Manji, 2000) and longitudinal studies have reported progressive increases in measures of grey matter volume following lithium initiation in bipolar disorder (Lyoo et al., 2010). These studies have generally shown that the effects of lithium on MRI estimates of grey matter volume are not uniform across the brain, and may localise to candidate regions such as the hippocampus, amygdala, anterior cingulate and prefrontal cortex (Hibar et al., 2017; Hibar et al., 2016; Monkul et al., 2007; Zung et al., 2016).

Whilst the effects of lithium on white matter are not fully understood, it has been consistently shown to positively influence measures of white matter integrity. Diffusion MRI (dMRI) enables the examination of the microstructural properties of tissue by mapping water molecule diffusion. In pure fluid states, water diffuses equally in all directions (isotropic). In neurons however, water cannot readily cross the hydrophobic myelin sheaths and diffusion is restricted (anisotropic) along the direction of the axon. The parallel configuration of axons in white matter renders this directional diffusion detectable at the microstructural scale, reflected in the dMRI-derived parameter of fractional anisotropy (FA). Decreases in FA within white matter have been shown to correspond with demyelination and axonal injury (Boretius et al., 2012; Song et al., 2002), such that FA is held to be a measure of white matter integrity. A number of studies have examined regional FA in bipolar disorder, generally reporting low FA values in white matter regions and tracts involved in mood regulation (Adler et al., 2006; Benedetti et al., 2011; Bruno, Cercignani, & Ron, 2008). The findings from whole brain dMRI studies are heterogeneous, but meta-analysis has identified two significant clusters of low FA in the right hemisphere (parahippocampal gyrus and the anterior cingulate/subgenual cingulate cortex) (Marlinge, Bellivier, & Houenou, 2014). White matter FA is greater in patients with bipolar disorder taking lithium compared to those on other medications, and the differences may be a function of the duration of treatment (Gildengers et al., 2015; Haarman et al., 2016; Macritchie et al., 2009). In a recent region of interest study using longitudinal dMRI data from adolescents with bipolar disorder (Kafantaris, Spritzer, Doshi, Saito, & Szeszko, 2017), FA values in the cingulum hippocampus were lower than healthy controls but increased following treatment with lithium, the effect being most marked in responders. However, in the absence of a whole brain analysis, it remains unclear whether this effect of lithium is regionally specific or global.

Regionally specific effects of lithium would indicate either that brain regions are differentially sensitive to its effects or that the drug itself is heterogeneously distributed across the brain. Recent advances in multinuclear MRI (^7^Li-MRI) have permitted the rapid determination of brain lithium distribution *in vivo* (Smith et al., 2018), demonstrating a heterogeneous distribution of lithium in bipolar disorder, in keeping with previous rodent imaging and human magnetic resonance spectroscopy studies (Lee et al., 2012; Zanni et al., 2017). It would therefore be reasonable to propose that the effects of lithium on white matter integrity relate to its regional concentration and the combination of ^7^Li-MRI and dMRI affords the capability to explore this.

In this study we compared generalised FA (gFA), a measure analogous to FA with the ability to account for crossing fibres (Tuch, 2004), across white matter regions of interest in patients with bipolar disorder receiving lithium, those taking other maintenance medications but naïve to lithium, and a healthy comparator group. Patients receiving lithium additionally underwent an assessment of brain lithium distribution by ^7^Li-MRI. We hypothesised that gFA values in patients receiving lithium treatment would exceed those of patients on other treatments and be comparable to healthy controls, and that the areas of the brain with the greatest signal intensity on ^7^Li-MRI would be those with the largest magnitude gFA in the lithium treated group.

## METHODS

### Subjects and assessments

Twenty-nine euthymic subjects with a diagnosis of bipolar disorder (I or II) and sixteen healthy control subjects recruited to the Bipolar Lithium Imaging and Spectroscopy Study (BLISS) were studied. Of those with bipolar disorder, twelve were taking lithium as a long-term treatment (Bipolar Disorder Lithium, BDL) and seventeen were taking other maintenance treatments but were naïve to lithium (Bipolar Disorder Control, BDC). The healthy control subjects (HC) had no history of psychiatric illness and were not taking any psychotropic medications. All subjects provided written informed consent and the study was granted a favourable ethical opinion by a United Kingdom National Research Ethics Committee (14/NE/1135). Exclusion criteria were contraindications to magnetic resonance examination, current or recent harmful drug or alcohol use, comorbid psychiatric diagnosis, learning disability, impairment of capacity or current liability to detention under the Mental Health Act 1983 (amended 2007). Diagnosis was confirmed through clinical interview using the NetSCID diagnostic tool (a validated online version of the Structured Clinical Interview for DSM-V Criteria; Telesage, Inc., Chapel Hill, NC, USA). All interviews and objective ratings were conducted by a trained research assistant (CJF) and discussed with a senior psychiatrist (DAC). Euthymic mood state was confirmed at entry to the study, defined as scores of less than seven on both the 21-item Hamilton Depression Rating Scale (HAM-D) and the Young Mania Rating Scale (YMRS). BDL subjects had taken lithium carbonate (Priadel™ modified release once daily) regularly for at least one year at the time of recruitment and all were taking at least one concomitant medication. All scans were performed at 9 am and the BDL subjects were instructed to take their lithium as usual the night before. Blood samples were taken immediately prior to scanning for confirmation that serum lithium levels fell within the target therapeutic range (0.6 – 1.0 mmol/L).

### Image acquisition

#### Scanner and coil systems

all MR data were acquired on a Philips 3T Achieva scanner (Philips Medical Systems, Best, The Netherlands) with ^1^H structural and diffusion weighted imaging performed using a Philips 8-channel SENSE head coil in all subjects. ^7^Li-MRI, together with an additional ^1^H structural image for the purposes of registration, was performed using a double tuned ^1^H/^7^Li radiofrequency (RF) birdcage head coil (RAPID Biomedical, Rimpar, Germany).

#### T1-weighted imaging acquisition

3D T1-weighted images (T1w) of brain anatomy, acquired in all subjects using the 8-channel SENSE head coil, were obtained with a ^1^H gradient echo sequence (TR = 9.6 ms, TE = 4.6 ms, FOV = 240 × 240 × 180 mm^3^, acquisition matrix = 240 × 208 × 180, acquisition voxel size = 1 × 1.15 × 1 mm^3^, reconstructed into a matrix size of 256 × 256 × 180, 1 average). Those in the BDL group were also scanned using a double tuned ^1^H/^7^Li head coil, with a ^1^H gradient echo sequence (TR = 8.2 ms, TE = 4.6 ms, FOV = 216 × 240 × 175 mm^3^, acquisition matrix = 180 × 200 × 146, acquisition voxel = 1.2 × 1.2 × 1.2 mm^3^ reconstructed into a matrix size of 240 × 240 × 146, 1 average). The total acquisition time for each sequence was less than 5 minutes.

#### Diffusion-weighted imaging acquisition

dMRI data was acquired using a single-shot pulsed gradient spin echo (EPI) sequence (TR = 10200 ms, TE = 74 ms). For each subject, 64 b1000 diffusion-weighted volumes were collected along non-collinear directions. In addition, one b0 volume was also acquired. Each volume consisted of 55 transverse slices (FOV = 240 × 240 × 137.5 mm^3^, voxel size = 2.5 mm isotropic, no gap). The total acquisition time for this sequence was less than 13 minutes.

#### ^7^Li magnetic resonance imaging

^7^Li 3D Balanced Steady State Free Precession (bSSFP) gradient echo acquisition (^7^Li-MRI) was acquired in eleven of the BDL subjects using a protocol constructed for maximal ^7^Li signal amplitude (FOV = 480 × 480 × 175 mm^3^, 20 × 19 × 7 acquisition matrix and 24.0 × 25.3 × 25.0 mm^3^ voxel size, with TR = 9.4 ms, TE = 4.5 ms, flip angle = 60 degrees, receiver bandwidth = 219 Hz/pixel, 500 averages per dynamic, 3 dynamics). Data were reconstructed into a 32 × 32 × 7 matrix with a voxel size of 15 × 15 × 25 mm^3^. The sequence, lasting approximately eight minutes, was conducted thrice without a notable interval and the three scan dynamics were averaged in complex form.

### Image processing and analysis

All T1w, dMRI and ^7^Li-MRI images were exported in DICOM format and converted to NIFTI format data using the Matlab (Mathworks^®^ Inc., Natick, MA, USA) toolbox ‘DICOM to NIfTI’ (Xiangrui, 2017). Data pre-processing and analysis were performed using Nipype, a Python based platform that provides a uniform interface to existing neuroimaging software and facilitates interaction between these packages within a single workflow (Gorgolewski et al., 2011). An outline of the processing pipeline is shown in Figure 1.

#### T1w image processing

T1w structural images acquired during the ^1^H imaging session were registered to the standard Montreal Neurological Institute (MNI) 2 mm brain using the FMRIB Linear and Non Linear Image Registration Tools (Jenkinson, Beckmann, Behrens, Woolrich, & Smith, 2012). These images were also linearly co-registered to the ^1^H-T1w image acquired during the ^7^Li imaging session using FMRIB’s Linear Image Registration Tool (Jenkinson et al., 2012). T1w structural images acquired during the ^7^Li imaging session were segmented into three tissue classes (grey matter, white matter and cerebrospinal fluid) using the FMRIB Automated Segmentation Tool (Zhang, Brady, & Smith, 2001).

#### Diffusion-weighted image processing

dMRI images from every subject were corrected for motion and eddy-current distortions using the eddy correct and bvec rotation utility in FSL (Andersson & Sotiropoulos, 2016). The b0 image was used as the reference image for realignment of all the dMRI images, and mean displacement for each subject across the acquisition was calculated. One-way ANOVA failed to reveal a significant difference between the mean displacement between the three groups (*p* = 0.64). Subsequently, skull-stripping was performed on the b0 image using FSL’s Brain Extraction Tool (Smith et al., 2002) and images were visually inspected to ensure skull-stripping had occurred successfully. A Constant Solid Angle (Q-Ball) model with fourth-order spherical harmonics was fitted to motion-corrected dMRI images using the Python package Dipy (version 0.11) (Garyfallidis et al., 2014) yielding an estimation of gFA for each brain voxel. gFA was chosen in favour of FA owing to the ability of the Q-Ball model to account for crossing fibres (Tuch, 2004). gFA maps were co-registered with structural T1w images using FMRIB’s Linear Image Registration Tool (Jenkinson et al., 2012) with 6 degrees of freedom. gFA maps were transformed into MNI space by applying the transformation matrix obtained from registering the structural T1w images to the standard MNI 2mm brain using FMRIB’s Non Linear Image Registration Tool (Andersson, Jenkinson, & Smith, 2007).

For the analysis of mean gFA across the white matter for each subject in MNI space, the mean gFA across all voxels in the John Hopkins University (JHU) White-Matter Tractography region of interest (ROI) atlas (Wakana, Jiang, Nagae-Poetscher, van Zijl, & Mori, 2004) was calculated. To quantify differences across groups (BDC, BDL, and HC), the effects of age and sex were regressed out and the residuals were compared between the groups. One-way ANOVA was used to test for group differences in mean white matter gFA residuals, followed by post-hoc t-tests (applying Tukey’s correction for multiple comparisons) to determine the direction of effect. Figure 1A shows a schematic outline of this processing step.

For a ROI based analysis of gFA in MNI space, region-wise differences in mean gFA were determined for each JHU ROI by computing effect sizes (Cohen’s d) between the groups. Effect sizes were computed using residuals after regressing out the effects of age and sex for each individual ROI.

#### ^7^Li-MRI image processing

^7^Li image analysis was performed in native space due to the relatively low spatial resolution of these images. After segmentation of the structural T1w image in subject native space, percentage tissue content (grey matter, white matter and cerebrospinal fluid) was estimated for each ^7^Li-MRI voxel. gFA maps were linearly registered to the same subject space and the relationship between mean gFA and lithium signal within each ^7^Li-MRI voxel was investigated at white matter thresholds ranging from 0-100% in steps of 25%. In essence, we performed all analyses steps at the resolution of the lithium images in subject space. This ensured that no effect of up-sampling, or interpolation of low resolution images would affect our results. Fig. 1B shows a schematic outline of this processing step.

To enable a group analysis of the association between gFA and ^7^Li-MRI signal intensity, a linear mixed effects model was used to account for inter-subject variations (Winter, 2013). As fixed effects, we entered gFA, age and sex (without interaction term) into the model. As random effects, we had intercepts for subjects and by-subject random slopes for the effect of ^7^Li-MRI signal. Visual inspection of residual plots did not reveal any obvious deviations from homoscedasticity or normality. P-values were obtained by likelihood ratio tests of the full model with the fixed effect of ^7^Li-MRI signal against the model without the fixed effect of ^7^Li-MRI signal. Supplementary Information A shows the full linear mixed effect analysis and additional tests to determine the random effects.

**Figure 1.**
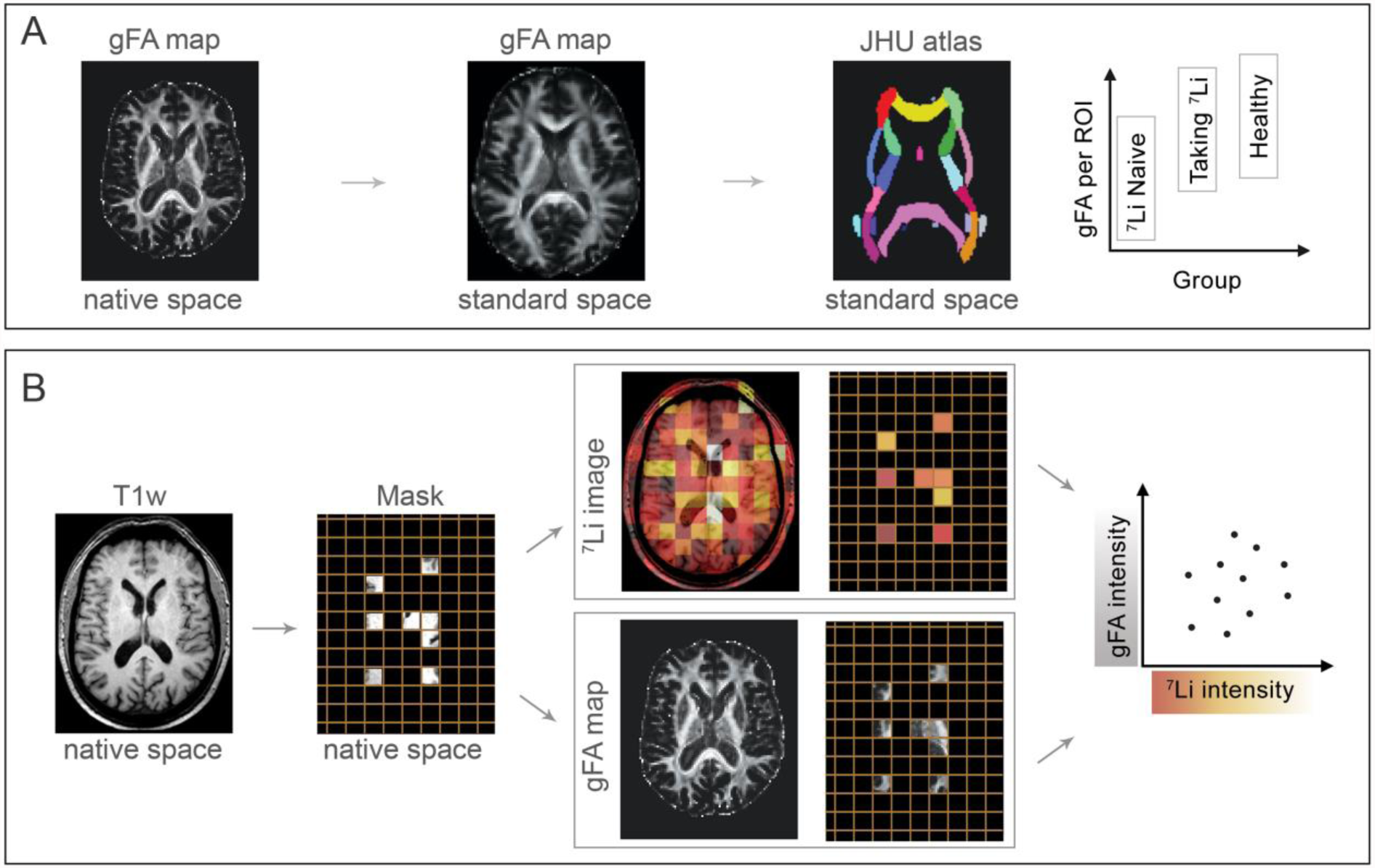
Schematic outline of processing pipeline. (A) gFA ROI analysis between groups was performed by transforming the gFA maps into standard space and applying the John Hopkins University (JHU) atlas. (B) The gFA ∼ lithium analysis for each subject was performed in native space at the resolution of the lithium image. The T1w image was used to identify lithium voxels containing a certain percentage of white matter, which essentially creates a mask. The mask was applied to both the gFA map as well as the ^7^Li-MRI image. Finally, mean gFA in each mask voxel was plotted against the ^7^Li signal. Hence, each data point corresponds to a lithium voxel in a subject.

## RESULTS

### Group characteristics

Twelve BDL subjects (five women; mean age: 46 ± 13 SD years), seventeen BDC subjects (eleven women; mean age: 44 ± 12 SD years) and sixteen HC subjects (ten women; mean age: 49 ± 3 SD years) were included in the analysis. One BDL subject was excluded from the ^7^Li-MRI analysis because their data was acquired during the ^7^Li-MRI sequence development phase and the acquisition protocol was inconsistent with the other subjects in the study. The groups did not differ in mean age (*p* = 0.77), sex distribution (*p* = 0.45) or educational level (*p* = 0.87) (Table 1). ANOVA followed by post-hoc testing of clinical information revealed that the HC group had significantly lower HAM-D scores than the BD group as a whole (*p* = 0.03).

**Table 1.** Subject characteristics. YMRS: Young Mania Rating Scale. HAM-D: Hamilton Rating Scale for Depression. LISERS: Lithium Side Effects Rating Scale. Values reported as mean (standard deviation).

|  | **Bipolar Disorder**<br>**Lithium**<br>**(n = 12)** | **Bipolar Disorder**<br>**Control**<br>**(n = 17)** | **Healthy**<br>**Control**<br>**(n = 16)** | **Significance** |
| --- | --- | --- | --- | --- |
| Sex (M/F) | 7/5 | 6/11 | 6/10 | p = 0.45 |
| Age (years) | 46(13) | 44(12) | 49(3) | p = 0.77 |
| Educational level (years) | 14(2) | 14(2) | 17(3) | p = 0.87 |
| YMRS score | 1(1) | 1(1) | 0.2(0.5) | p = 0.96 |
| HAM-D score | 5(6) | 5(5) | 1(1) | p = 0.03 |
| Duration of lithium treatment (years) | 8(9) | n/a | n/a | n/a |
| Priadel™ dose (mg) | 900(130) | n/a | n/a | n/a |
| Serum lithium concentration (mmol/L) | 0.8(0.13) | n/a | n/a | n/a |
| LISERS score | 17(13) | n/a | n/a | n/a |

All subjects with bipolar disorder were taking some form of medication at time of scanning. Fisher’s exact tests were performed in order to compare the use of medication class between diagnostic groups. Barring lithium, no significant differences were found between the bipolar disorder groups across all major medication classes. Medication usage information for each group and their corresponding *p* values are provided in Table 2.

**Table 2.** Medication use by class. Fisher’s exact tests found no differences in medication class between diagnostic groups other than lithium (OR: Odds Ratio).

| Medication class | Bipolar Disorder<br>Lithium (n=12) | Bipolar Disorder<br>controls (n=17) | Significance |
| --- | --- | --- | --- |
| Antipsychotics | 8 (66%) | 11 (65%) | OR = 1.1, p = 1 |
| Antidepressants | 6 (50%) | 10 (59%) | OR = 0.7, p = 0.7 |
| Anticonvulsants | 5 (42%) | 10 (59%) | OR = 0.5, p = 0.5 |
| Anxiolytics | 3 (25%) | 4 (24%) | OR = 1.1, p = 1 |
| Hypnotic | 3 (25%) | 2 (12%) | OR = 2.5, p = 0.6 |
| Antihistamine | 0 | 1 (6%) | OR = 0, p = 1 |
| Over the counter | 0 | 1 (6%) | OR = 0, p = 1 |
| Mood stabilisers / mania | 12 (Lithium, 100%) | 0 | n/a |

### Mean white matter gFA comparison between groups

A significant negative correlation (r = −0.52; *p* = 0.0002) between mean white matter gFA and age was found across all subjects. This was anticipated as FA has previously been shown to be negatively associated with age in adulthood (Lebel, Caverhill-Godkewitsch, & Beaulieu, 2010). After regressing out the effects of age and sex, ANOVA revealed a significant difference between the groups (F(2, 42); *p* = 0.02; Figure 2b) and post hoc tests revealed that the BDL and HC groups exhibited higher mean white matter gFA residuals compared to the BDC group (t = 2.5, *p* = 0.05; t = 2.7, *p* = 0.03, respectively). No differences were found between the BDL and HC groups (t = 0.02 *p* = 1).

**Figure 2.**
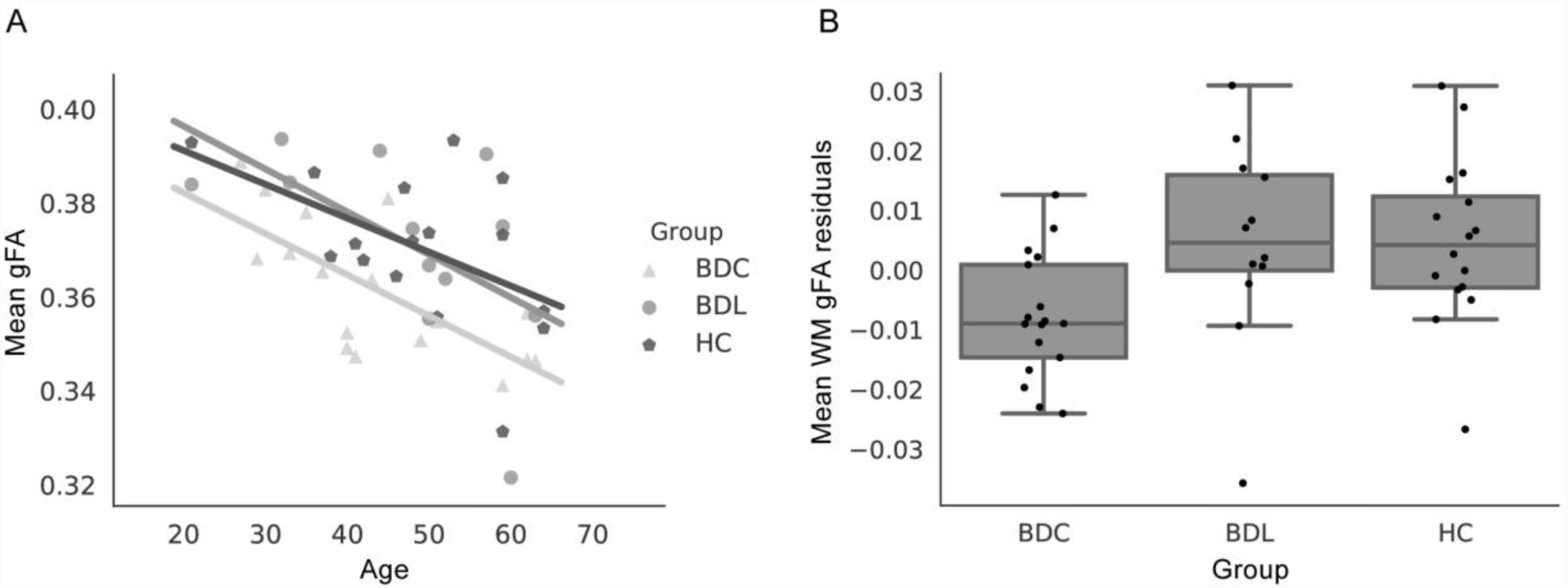
gFA group comparison. (A) Association between mean white matter gFA and age. Each dot is an individual subject and lines indicate the least squared regressions. (B) Group comparison of mean gFA residuals after correction for age and sex.

### Region of interest gFA group comparison

Region-wise gFA effect sizes for the comparison BDL > BDC are shown in Figure 3. Subjects taking lithium exhibited higher gFA in 44 out of 48 ROIs compared to those taking other medications for bipolar disorder but naïve to lithium, with eight regions exhibiting an effect size of greater than 0.8 (signifying a large effect). Effect sizes ranged from −0.3 to + 1.5, indicating spatial heterogeneity in gFA differences between the groups. All ROI labels and their corresponding effect sizes are provided in Supplementary Information B.

**Figure 3.**
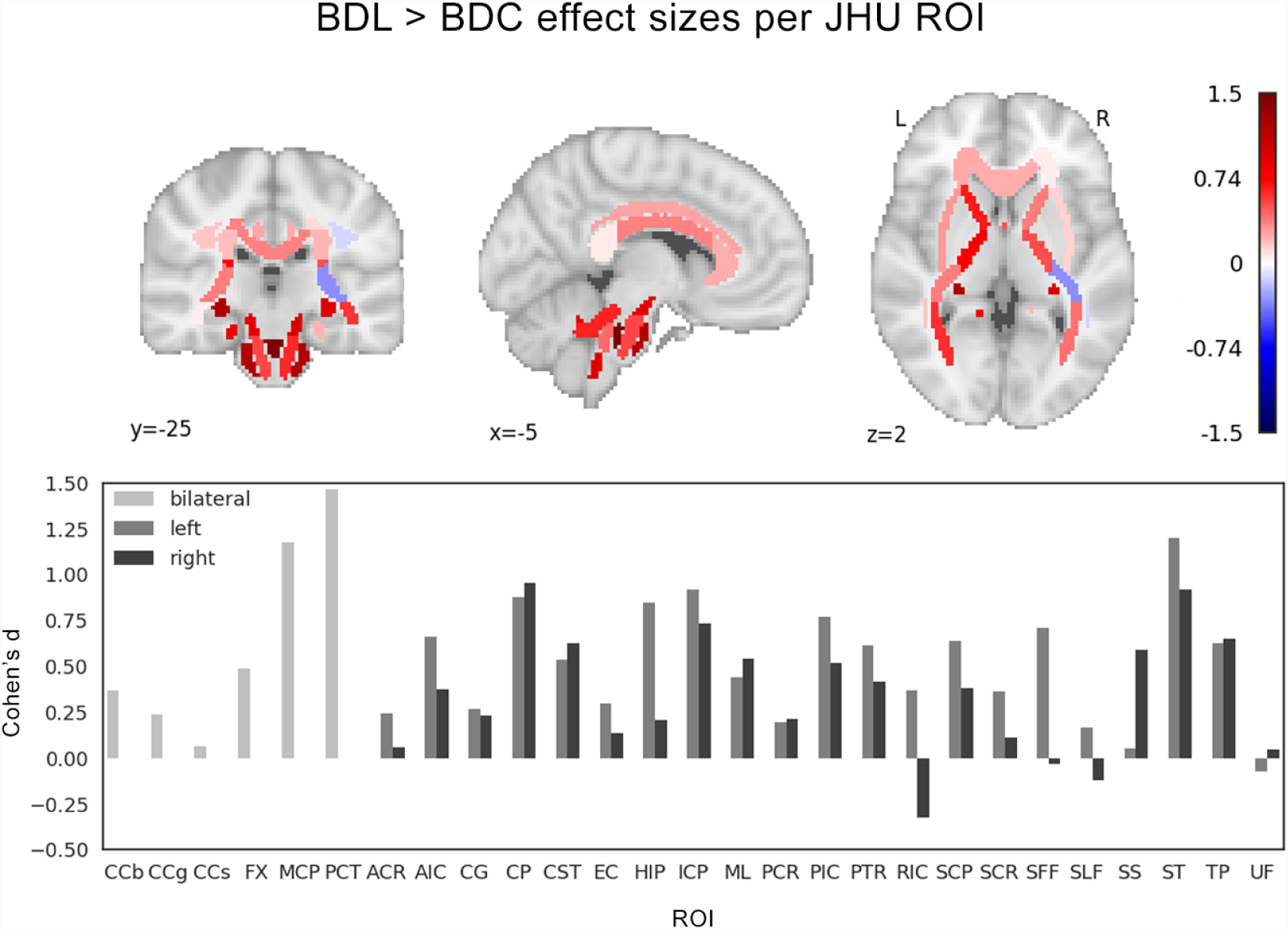
Region-wise gFA effect sizes (Cohen’s d) per JHU ROI comparing BDL>BDC. Top: JHU ROIs in standard space are colour coded according to the effect size of BDL>BDC. Bottom: Bar plot showing individual effect sizes for each ROI. Full ROI labels and all effect sizes are provided in Supplementary B. Note effect sizes are calculated on residual gFA values after age and sex correction for each ROI.

### Co-localisation of lithium and gFA

The relationship between white matter integrity and the spatial distribution of lithium was determined in a linear mixed effect analysis of ^7^Li-MRI signal intensity and gFA values in voxels containing varying proportions of white matter at the resolution of the ^7^Li-MRI scan (Figure 4). Results revealed a highly significant association between the ^7^Li-MRI signal and gFA (*p* < 0.01) once white matter content exceeded 50% in the ^7^Li-MRI voxel of interest.

Using linear mixed effect modelling, we also tested the random slope model against a random intercept model to determine whether there was evidence to suggest that individual subjects need to be modelled with individual slopes. These results revealed there was a significant difference in the models accounting for a subject specific slope versus one that does not (*p* < 0.05). The full linear mixed effect analysis is described in detail in Supplementary Information A.

**Figure 4.**
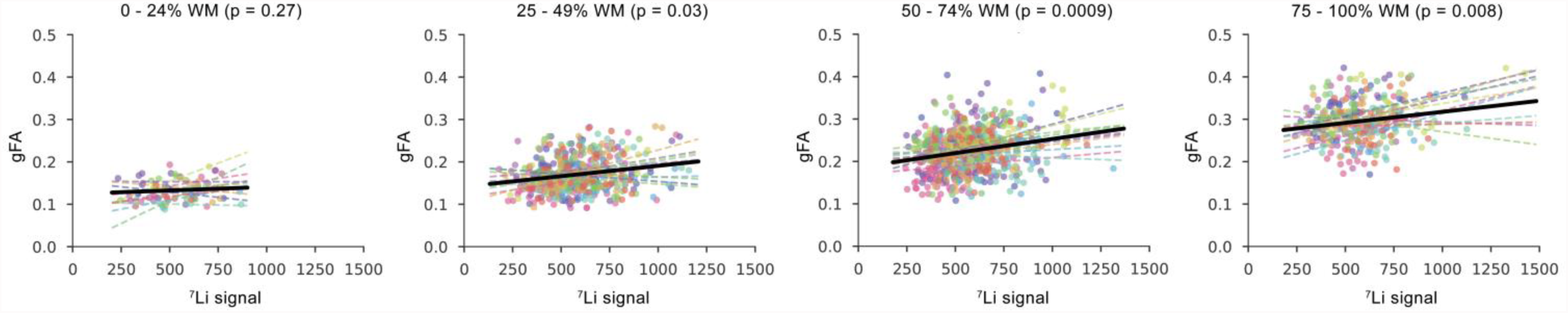
Relationship between lithium signal amplitude and gFA. Relationship between ^7^Li-MRI signal intensity and mean gFA in ^7^Li-MRI voxels containing varying levels of white matter (ranging from 0-100% in steps of 25%). Dashed lines represent individual subject least-squares regression line. Black line represents group linear mixed effects regression line. P values represent significance of a theoretical likelihood test comparing linear mixed effect models which do and do not include ^7^Li signal as a fixed effect.

## DISCUSSION

In this study, we report that patients with bipolar disorder treated with lithium have white matter gFA values comparable to healthy controls and greater than bipolar disorder patients receiving other medications. Further, as hypothesised, the spatial distribution of the ^7^Li-MRI signal positively correlated with gFA. Our findings add to the growing literature suggesting lithium that use is associated with a rectification of the microstructural MRI abnormalities that occur in bipolar disorder (Hafeman et al., 2012; Hibar et al., 2017; Hibar et al., 2016; Kafantaris et al., 2017; Moore et al., 2000).

One major challenge in our work was the relatively low resolution of the lithium image compared to the T1w and dMRI images, meaning that a single lithium voxel often contained mixed tissue types. To enable a fair comparison, we performed all analysis using the lithium signal in native space at the resolution of the lithium image. Our results from linear mixed effect modelling show the emergence of a robust and highly significant effect (*p* < 0.01) of the lithium signal in explaining gFA in lithium voxels containing over 50% of white matter. Furthermore, we found a significant difference between the random intercept model and the random slope model, potentially suggesting subject-specific slopes of the gFA ˜ lithium relationship. However, these results reached statistical power through the use of linear mixed effect modelling by estimating associations on the BDL group as a whole. Our data did not permit us to derive reliable subject specific slopes. Hence, at the present stage, we cannot conclude the significance of individual slopes, nor can we infer the causality of the association between gFA and lithium.

Lithium is purported to exert a range of neurorestorative effects (Forlenza, De-Paula, & Diniz, 2014; Quiroz, Rodrigo, Zarate, Jr, & Manji, 2010), many of which could influence white matter microstructure and affect gFA. For instance, lithium is known to inhibit glycogen synthase kinase 3β and inhibition of this enzyme increases the number of oligodendrocytes and promotes peripheral nerve myelination in mice (Azim & Butt, 2011; Makoukji et al., 2012). Lithium treatment has been shown to be positively associated with MRI measures of white matter integrity (Gildengers et al., 2015; Haarman et al., 2016; Macritchie et al., 2009) as well as protect against a decrease in white matter volume over time (Berk et al., 2017). Lithium has been associated with an increase in FA in the cingulum hippocampus in a recent longitudinal study of adolescents with bipolar disorder, where responders to treatment showed the greatest percentage increase in FA (Kafantaris et al., 2017).

Lithium related alterations in water homeostasis also require consideration in the interpretation of its effects on dMRI-derived measures. Rats fed lithium for five weeks have elevated brain tissue water content (Phatak, Shaldivin, King, Shapiro, & Regenold, 2006) and it has been argued that this may underpin the volumetric changes detected in neuroimaging studies in bipolar disorder (Regenold, 2008). Phatak et al., 2006 reported that water content was significantly increased only in the frontal cortex, suggesting regional specificity of action but not accounting for the more widespread cortical differences in grey matter volume on MRI. However, increased water content driven by lithium cannot easily explain its effects on FA, as widespread increases in free water content within white matter has been linked to decreased FA in first-episode patients with psychosis (Lyall et al., 2017).

In practice, lithium may be preferentially prescribed to those deemed more likely to respond to it, to those with more severe presentations or when there is confidence that the patient will manage regular blood level monitoring. It could be argued that the differences in FA between patients treated with lithium and those on other medications reflects an as yet unidentified illness characteristic driving a selection bias rather than a direct action of lithium. However, longitudinal data (Kafantaris et al., 2017) and our finding of a relationship between gFA and regional lithium concentration strongly suggests that lithium is directly involved.

Further, the effects of lithium on white matter may be linked to its clinical efficacy, holding the potential to predict therapeutic response (Berk et al., 2017; Kafantaris et al., 2017). In an elderly population with bipolar disorder, a longer duration of lithium treatment was found to be associated with higher FA but interestingly, not with better cognitive performance (Gildengers et al., 2015). Investigating patients with bipolar disorder initiating lithium for the treatment of depression, Machado-Vieira et al. (2016) reported that serum and brain lithium concentration only correlated in those who entered remission, but their ^7^Li magnetic resonance spectroscopy sequence did not permit regional localisation. The potential to use changes in FA as a predictor of therapeutic response may be enhanced by being able to co-localise FA with the spatial distribution of lithium. In future work, multimodal imaging studies utilising ^7^Li-MRI and larger cohorts that permit the segregation of subjects into groups based upon lithium response may reveal localised response-specific relationships between regional brain lithium concentration and MRI derived measures such as gFA. Such approaches may help characterise lithium responders and move us a step closer to personalised medicine in bipolar disorder.

In conclusion, we report dMRI results that are consistent with the previous literature and a novel observation of the correlation between gFA and regional lithium concentration. ^7^Li-MRI is a new technique that holds potential to provide multiple insights into the effects of lithium in the brain in bipolar disorder. However, the limited spatial resolution of this technique restricted the present study to reporting results based on voxels containing mixed tissue types. Despite this, we found a consistent significant relationship between ^7^Li-MRI signal and gFA in voxels containing more than 50% white matter. Whilst challenging, there is scope for on-going technical development of ^7^Li-MRI with clear potential to improve the spatial resolution of lithium images. The ability to co-localise a psychotropic medication with its effect *in vivo* spurs us to meet these challenges.

## Acknowledgements

We gratefully acknowledge our funding sources, the Medical Research Council of the United Kingdom (Clinician Scientist Fellowship BH135495 to DAC), the Wellcome Trust (Seed award to YW 208940/Z/17/Z and Seed award to PNT 210109/Z/18/Z), and the Reece Foundation (PhD Studentship to JN). We thank Dr Matthew Clemence (Philips Medical Science) for technical expertise and guidance with multinuclear MR.

Data included in this manuscript was previously presented in abstract form (Biological Psychiatry May 1 2018; Volume 83 (9), Supplement pages S99-100), and will be submitted as a pre-print to bioRxiv.org.

## References

Adler, C. M., Adams, J., DelBello, M. P., Holland, S. K., Schmithorst, V., Levine, A.,…Strakowski, S. M. (2006). Evidence of white matter pathology in bipolar disorder adolescents experiencing their first episode of mania: A diffusion tensor imaging study. American Journal of Psychiatry, 163(2), 322–324.

Andersson, Jenkinson, & Smith. (2007). Non-linear registration, aka Spatial normalisation FMRIB technical report TR07JA2.

Andersson, & Sotiropoulos. (2016). An integrated approach to correction for off-resonance effects and subject movement in diffusion MR imaging. Neuroimage, 125, 1063–1078. doi:10.1016/j.neuroimage.2015.10.019

Azim, K., & Butt, A. M. (2011). GSK3β negatively regulates oligodendrocyte differentiation and myelination in vivo. Glia, 59(4), 540–553. doi:10.1002/glia.21122

Benedetti, F., Yeh, P.-H., Bellani, M., Radaelli, D., Nicoletti, M. A., Poletti, S.,… Brambilla, P. (2011). Disruption of white matter integrity in bipolar depression as a possible structural marker of illness. Biol. Psychiatry, 69(4), 309–317. doi:10.1016/j.biopsych.2010.07.028

Berk, M., Dandash, O., Daglas, R., Cotton, S. M., Allott, K., Fornito, A.,… Yücel, M. (2017). Neuroprotection after a first episode of mania: a randomized controlled maintenance trial comparing the effects of lithium and quetiapine on grey and white matter volume. Transl. Psychiatry, 7(2), e1041. doi:10.1038/tp.2017.13

Boretius, S., Escher, A., Dallenga, T., Wrzos, C., Tammer, R., Brück, W.,… Stadelmann, C. (2012). Assessment of lesion pathology in a new animal model of MS by multiparametric MRI and DTI. Neuroimage, 59(3), 2678–2688. doi:10.1016/j.neuroimage.2011.08.051

Bruno, S., Cercignani, M., & Ron, M. A. (2008). White matter abnormalities in bipolar disorder: a voxel-based diffusion tensor imaging study. Bipolar Disord, 10(4), 460–468.

Forlenza, O. V., De-Paula, V. J. R., & Diniz, B. S. O. (2014). Neuroprotective Effects of Lithium: Implications for the Treatment of Alzheimer’s Disease and Related Neurodegenerative Disorders. ACS Chem. Neurosci., 5(6), 443–450. doi:10.1021/cn5000309

Garyfallidis, E., Brett, M., Amirbekian, B., Rokem, A., van der Walt, S., Descoteaux, M.,… Dipy, C. (2014). Dipy, a library for the analysis of diffusion MRI data. Front. Neuroinform., 8, 8. doi:10.3389/fninf.2014.00008

Gildengers, A. G., Butters, M. A., Aizenstein, H. J., Marron, M. M., Emanuel, J., Anderson, S. J.,… Reynolds, C. F. (2015). Longer lithium exposure is associated with better white matter integrity in older adults with bipolar disorder. Bipolar Disorders, 17(3), 248–256. doi:10.1111/bdi.12260

Gorgolewski, K., Burns, C. D., Madison, C., Clark, D., Halchenko, Y. O., Waskom, M. L., & Ghosh, S. S. (2011). Nipype: a flexible, lightweight and extensible neuroimaging data processing framework in python. Front. Neuroinform., 5, 13. doi:10.3389/fninf.2011.00013

Haarman, B. C. M., Riemersma-Van der Lek, R. F., Burger, H., de Groot, J. C., Drexhage, H. A., Nolen, W. A., & Cerliani, L. (2016). Diffusion tensor imaging in euthymic bipolar disorder - A tract-based spatial statistics study. J Affect Disord, 203, 281–291. doi:10.1016/j.jad.2016.05.040

Hafeman, D. M., Chang, K. D., Garrett, A. S., Sanders, E. M., & Phillips, M. L. (2012). Effects of medication on neuroimaging findings in bipolar disorder: an updated review. Bipolar Disord., 14(4), 375–410. doi:10.1111/j.1399-5618.2012.01023.x

Hibar, D. P., Westlye, L. T., Doan, N. T., Jahanshad, N., Cheung, J. W., Ching, C. R. K.,… Andreassen, O. A. (2017). Cortical abnormalities in bipolar disorder: an MRI analysis of 6503 individuals from the ENIGMA Bipolar Disorder Working Group. Mol. Psychiatry. doi:10.1038/mp.2017.73

Hibar, D. P., Westlye, L. T., van Erp, T. G. M., Rasmussen, J., Leonardo, C. D., Faskowitz, J.,… Andreassen, O. A. (2016). Subcortical volumetric abnormalities in bipolar disorder. Mol. Psychiatry, 21(12), 1710–1716. doi:10.1038/mp.2015.227

Jenkinson, M., Beckmann, C. F., Behrens, T. E. J., Woolrich, M. W., & Smith, S. M. (2012). FSL. Neuroimage, 62(2), 782–790. doi:10.1016/j.neuroimage.2011.09.015

Kafantaris, V., Spritzer, L., Doshi, V., Saito, E., & Szeszko, P. R. (2017). Changes in white matter microstructure predict lithium response in adolescents with bipolar disorder. Bipolar Disord, 19(7), 587–594. doi:10.1111/bdi.12544

Lebel, C., Caverhill-Godkewitsch, S., & Beaulieu, C. (2010). Age-related regional variations of the corpus callosum identified by diffusion tensor tractography. Neuroimage, 52(1), 20–31. doi:http://dx.doi.org/10.1016/j.neuroimage.2010.03.072

Lee, J.-H., Adler, C., Norris, M., Chu, W.-J., Fugate, E. M., Strakowski, S. M., & Komoroski, R. A. (2012). 4-T ^7^Li 3D MR spectroscopy imaging in the brains of bipolar disorder subjects. Magnetic Resonance in Medicine, 68(2), 363–368. doi:10.1002/mrm.24361

Lyall, A. E., Pasternak, O., Robinson, D. G., Newell, D., Trampush, J. W., Gallego, J. A.,… Szeszko, P. R. (2017). Greater extracellular free-water in first-episode psychosis predicts better neurocognitive functioning. Mol Psychiatry. doi:10.1038/mp.2017.43

Lyoo, I. K., Dager, S. R., Kim, J. E., Yoon, S. J., Friedman, S. D., Dunner, D. L., & Renshaw, P. F. (2010). Lithium-Induced Gray Matter Volume Increase As a Neural Correlate of Treatment Response in Bipolar Disorder: A Longitudinal Brain Imaging Study. Neuropsychopharmacology, 35(8), 1743–1750. doi:10.1038/npp.2010.41

Machado-Vieira, R., Otaduy, M. C., Zanetti, M. V., De Sousa, R. T., Dias, V. V., Leite, C. C.,… Gattaz, W. F. (2016). A Selective Association between Central and Peripheral Lithium Levels in Remitters in Bipolar Depression: A 3T-Li-7 Magnetic Resonance Spectroscopy Study. Acta Psychiatrica Scandinavica, 133(3), 214–220. doi:10.1111/acps.12511

Macritchie, Lloyd, A. J., Bastin, M. E., Vasudev, K., Gallagher, P., Eyre, R.,… Young, A. H. (2009). White matter microstructural abnormalities in euthymic bipolar disorder. Br. J. Psychiatry, 196(1), 52–58. doi:10.1192/bjp.bp.108.058586

Makoukji, J., Belle, M., Meffre, D., Stassart, R., Grenier, J., Shackleford, G. g.,… Massaad, C. (2012). Lithium enhances remyelination of peripheral nerves. Proc. Natl. Acad. Sci. U. S. A., 109(10), 3973–3978. doi:10.1073/pnas.1121367109

Marlinge, E., Bellivier, F., & Houenou, J. (2014). White matter alterations in bipolar disorder: potential for drug discovery and development. Bipolar Disorders, 16(2), 97–112. doi:10.1111/bdi.12135

Monkul, E. S., Matsuo, K., Nicoletti, M. A., Dierschke, N., Hatch, J. P., Dalwani, M.,… Soares, J. C. (2007). Prefrontal gray matter increases in healthy individuals after lithium treatment: A voxel-based morphometry study. Neuroscience Letters, 429, 7–11. doi:10.1016/j.neulet.2007.09.074|ISSN 0304-3940

Moore, G. J., Bebchuk, J. M., Wilds, I. B., Chen, G., & Manji, H. K. (2000). Lithium-induced increase in human brain grey matter.[erratum appears in Lancet 2000 Dec 16;356(9247):2104 Note: Menji HK [corrected to Manji HK]]. Lancet, 356(9237), 1241–1242.

Phatak, P., Shaldivin, A., King, L. S., Shapiro, P., & Regenold, W. T. (2006). Lithium and inositol: effects on brain water homeostasis in the rat. Psychopharmacology, 186(1), 41–47. doi:10.1007/s00213-006-0354-y

Quiroz, J. A., Rodrigo, M.-V., Zarate, C. A., Jr, & Manji, H. K. (2010). Novel Insights into Lithium’s Mechanism of Action: Neurotrophic and Neuroprotective Effects. Neuropsychobiology, 62(1), 50–60. doi:10.1159/000314310

Regenold, W. T. (2008). Lithium and increased hippocampal volume - More tissue or more water? Neuropsychopharmacology, 33(7), 1773–1774. doi:10.1038/sj.npp.1301524

Smith, Thelwall, P. T., Necus, J., Flowers, C. J., Blamire, A. M., & Cousins, D. A. (2018). 3D 7Li magnetic resonance imaging of brain lithium distribution in bipolar disorder. Molecular Psychiatry. doi:10.1038/s41380-018-0016-6

Smith, Zhang, Y. Y., Jenkinson, M., Chen, J., Matthews, P. M., Federico, A., & De Stefano, N. (2002). Accurate, robust, and automated longitudinal and cross-sectional brain change analysis. Neuroimage, 17(1), 479–489. doi:10.1006/nimg.2002.1040

Song, S.-K., Sun, S.-W., Ramsbottom, M. J., Chang, C., Russell, J., & Cross, A. H. (2002). Dysmyelination revealed through MRI as increased radial (but unchanged axial) diffusion of water. Neuroimage, 17(3), 1429–1436.

Tondo, L., Vazquez, G. H., & Baldessarini, R. J. (2014). Options for pharmacological treatment of refractory bipolar depression. Current Psychiatry Reports, 16(2), 431–431. doi:10.1007/s11920-013-0431-y

Tuch, D. S. (2004). Q-ball imaging. Magnetic Resonance in Medicine, 52(6), 1358–1372. doi:10.1002/mrm.20279

Wakana, S., Jiang, H., Nagae-Poetscher, L. M., van Zijl, P. C. M., & Mori, S. (2004). Fiber Tract–based Atlas of Human White Matter Anatomy. Radiology, 230(1), 77–87. doi:10.1148/radiol.2301021640

Winter. (2013). Linear models and linear mixed effects models in {R} with linguistic applications. CoRR, abs/1308.5499. doi:href="http://arxiv.org/abs/1308.5499

Xiangrui, L. (2017). DICOM to NIfTI converter, NIfTI tool and viewer version 2017.10.31. MATLAB Central File Exchange.

Zanni, G., Michno, W., Di Martino, E., Tjarnlund-Wolf, A., Pettersson, J., Mason, C. E.,… Hanrieder, J. (2017). Lithium accumulates in neurogenic brain regions as revealed by high resolution ion imaging. Scientific Reports, 7, 40726. doi:10.1038/srep40726

Zhang, Y., Brady, M., & Smith, S. (2001). Segmentation of brain MR images through a hidden Markov random field model and the expectation-maximization algorithm. Ieee Transactions on Medical Imaging, 20(1), 45–57. doi:10.1109/42.906424

Zung, S., Souza-Duran, F. L., Soeiro-de-Souza, M. G., Uchida, R., Bottino, C. M., Busatto, G. F., & Vallada, H. (2016). The influence of lithium on hippocampal volume in elderly bipolar patients: a study using voxel-based morphometry. Translational Psychiatry, 6, e846. doi:10.1038/tp.2016.97

